# Viral community diversity in the rhizosphere of the foundation salt marsh plant *Spartina alterniflora*

**DOI:** 10.1101/2025.04.03.647062

**Authors:** Isabelle Du Plessis, Hannah Snyder, Rachel Calder, José L. Rolando, Joel E. Kostka, Joshua S. Weitz, Marian Dominguez-Mirazo

## Abstract

Viruses of microorganisms impact microbial population dynamics, community structure, nutrient cycling, gene transfer, and genomic innovation. In wetlands, root-associated microbial communities mediate key biogeochemical processes important for plants involved in ecosystem maintenance. Nonetheless, the presence and role of microbial viruses in salt marshes remains poorly understood. In this study, we analyzed 24 metagenomes retrieved from the root zone of *Spartina alterniflora*, a foundation plant in salt marshes of the eastern and Gulf coasts of the U.S. The samples span three plant compartments—bulk sediment, rhizosphere, and root—and two cordgrass plant phenotypes: short and tall. We observed differentiation between phenotypes and increased similarity in viral communities between the root and rhizosphere, indicating that plant compartment and phenotype shape viral community composition. The majority of viral populations characterized are novel at the genus level, with a subset predicted to target microorganisms known to carry out key biogeochemical functions. The findings provide a holistic assessment of plant-associated viral diversity and community composition as well as identifying potential targets for exploring viral modulation of microbially-mediated ecosystem functioning in intertidal wetlands.

**Importance:** Salt marshes are vital coastal ecosystems. Microbes in these environments drive nutrient cycling and support plant health, with *Spartina alterniflora* serving as a foundation species. This study explores viral communities associated with *S. alterniflora*, revealing how plant compartment and phenotype shape viral composition. The discovery of numerous novel viruses, some potentially influencing microbes involved in key biogeochemical processes, highlights their ecological significance. Given the increasing pressures on coastal ecosystems, understanding virus-microbe-plant interactions is essential for predicting and managing ecosystem responses to environmental change.

## 1 Introduction

Salt marshes are intertidal wetland ecosystems that provide critical ecosystem services to coastal communities such as protection from storm surge, erosion control, carbon sequestration, and a supply of raw materials and food [1]. Smooth cordgrass, *Spartina alterniflora*, dominates the North American salt marshes on the Atlantic and Gulf of Mexico coastlines of North America. *S. alterniflora* is a foundation species that engineers the ecosystem through soil stabilization, building up the marsh platform, and stimulating the recycling of carbon and nutrients within the ecosystem [2]. Interactions between microbial populations and *S. alterniflora* plants facilitate biogeochemical activity in salt marshes, especially in the root zone, where plant microbiomes play a key role in coupling the sulfur cycle with both the carbon and nitrogen cycles [3, 4].

Viruses of microbes are highly abundant, diverse, and influence microbial community dynamics [5, 6]. The infection of microbes by viruses impacts ecosystem functioning by shifting microbial community dynamics [6, 7] and through direct metabolic modulation of infected cells (i.e., ‘virocells’) [7–10]. Viral community composition varies across ecosystems and environmental gradients including soil depth, chemical concentrations, and plant and microbial community composition [11–15]. Viruses of microbes are particularly abundant in soils, with typical virus-like particle estimates exceeding 10^9^ per g [16]. Furthermore, metagenomic studies of soils and wetlands have revealed that viral populations are exceptionally diverse, yet only a fraction of this diversity (and its putative impacts) has been characterized [11, 13].

Here, we describe the viral populations associated with *S. alterniflora* in Sapelo Island, Georgia, USA (a protected barrier island), across three microbiome compartments — bulk sediment, rhizosphere, and root — and two plant phenotypes, tall and short. Proximity to large tidal creeks impacts plant traits, with plants located closer to tidal banks exhibiting a tall phenotype while plants growing farther away exhibit a short phenotype [17]. The biomass gradient from tall to short *S. alterniflora* has been attributed to the interacting effects of sulfide toxicity, salinity stress, and anoxia, which more severely affect the short phenotype [17]. Prior analysis of bacterial metagenomes at this site revealed a predominance of sulfur-oxidizing and sulfur-reducing bacteria in *S. alterniflora* root microbiome, with root compartment and phenotype driving microbial diversity [3]. These microbes highly express genes involved in sulfur oxidation, carbon fixation, and nitrogen fixation, with higher levels of sulfur oxidation gene transcription linked to the stressed short phenotype [4]. Our analysis of viral sequences in bulk metagenomes reveals that plant phenotype and compartment are key drivers of viral community composition. We taxonomically describe the viral populations and find that most are novel at the genus level, while some are affiliated with genera previously identified in metagenomic studies of aquatic and soil-associated viral communities. Finally, we predict virus-host interactions and find that the viral populations associated with the *S. alterniflora* root zone target microbial orders that encompass sulfur-oxidizing, sulfate-reducing, and iron-oxidizing microbes. These findings expand our understanding of plant-associated viral communities and host-phage interactions in salt marsh environments.

## 2 Results

### 2.1 Relationship Between Viral Communities, Plant Phenotype, and Microbiome Compartment in *S. alterniflora*

Samples were collected in a previous study [4] from the *S. alterniflora* rhizosphere along established ∼100 m transects extending from a large tidal creek at GCE 06 within the NSF-sponsored Georgia Coastal Ecosystems Long-term Ecological Research site on Sapelo Island, Georgia, USA, during the summers of 2018 and 2019. Samples were collected from three different microbiome compartments, root, rhizosphere, and bulk sediment for two different plant phenotypes, short (*<* 50cm) and tall (*>* 80 cm), (Figure 1a). Bulk DNA was previously extracted and sequenced from each sample for metagenomic analysis of the plant-associated microbiome [4]. Here, we repurpose the sequencing data to characterize the viral populations associated with *S. alterniflora*.

**Figure 1:**
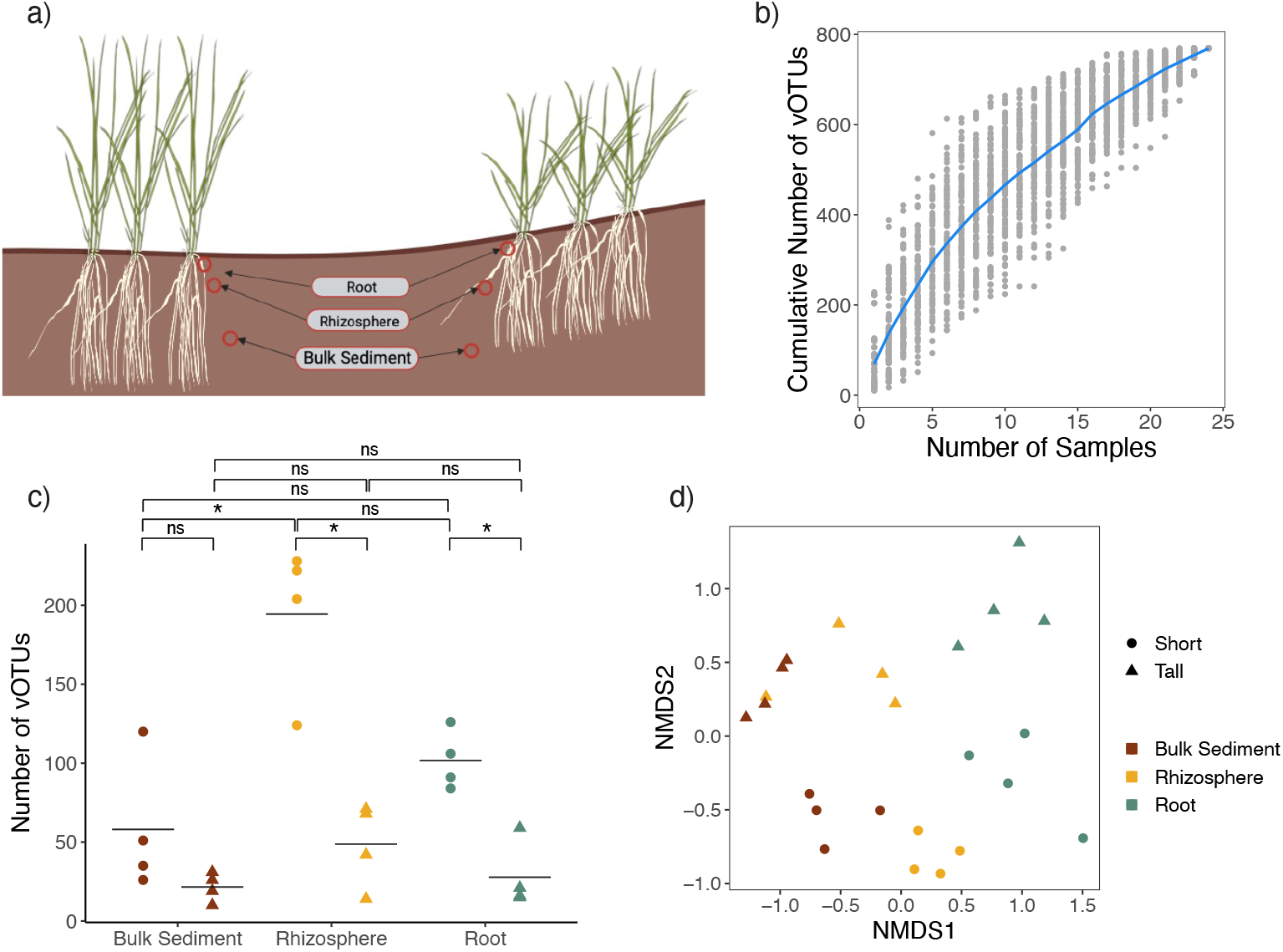
Viral communities in the *S. alterniflora* root zone are shaped by plant phenotype and microbiome compartment. **(a)** Metagenomes were obtained from the *S. alterniflora* rhizosphere at established transects on Sapelo Island, Georgia from three compartments (root, rhizosphere, and bulk sediment) and two plant phenotypes (short *<* 50 cm and tall *>* 80 cm), for a total of 24 samples. **(b)** Rarefection curve displaying the cumulative number of vOTUs found with increasing sample number, 100 permutations of sample order were performed (gray points). The mean cumulative number of vOTUs for each number of samples is represented by the blue line. The saturation plot shows that viral richness is not fully represented by the sampling and viral recovery efforts. **(c)** Viral richness, depicted by the number of vOTUs per phenotype per compartment. Horizontal lines represent the mean number of vOTUs among samples obtained from the same plant compartment and phenotype combination. Asterisks indicate statistical differences based on pairwise Mann–Whitney tests (two-sided, p-value *<* 0.05). **(d)** Non-metric multidimensional scaling (NMDS) of Bray-Curtis dissimilarity across each sample based on vOTU relative abundance (TPM) per sample. The NMDS shows similarities between viral population abundance among the samples, revealing separation between compartments and plant phenotypes. NMDS stress: 0.11.

Contigs obtained from *de novo* assembly were input into the viral identification tool Virsorter2 [18], dereplicated, and checked for quality (see Methods). For all samples combined, we recovered 769 viral operational taxonomic units (vOTUs) with sequences larger than 5kbp, of which 765 had not been previously described. The other 4 sequences were identified by read mapping to soil and aquatic viral databases [14, 19] (see Methods). The absence of saturation in the resulting rarefaction curve indicates that our sampling efforts did not fully capture the viral diversity within the system (Figure 1b). This is consistent with the fact that soils are highly diverse, and that bulk metagenomes are often ineffective for identifying viral populations in soil [20]. Viral-like particle (VLP) extraction and enrichment have proven to be more effective strategies for this purpose – an issue we revisit in the Discussion.

The presence of each vOTU in a given sample was assessed by read mapping. A vOTU was considered to be present in a sample if mapping resulted in ≥ 75% of the contig length covered. Viral richness is shown in Figure 1c. Samples from short plant phenotypes show higher viral richness than their tall counterparts. This observation is statistically significant for the rhizosphere, and root compartments only, with no significant difference in viral richness between short and tall samples of the bulk sediment. The viral populations recovered from the rhizosphere are more diverse than the bulk sediment for samples obtained from short phenotypes only.

The relative abundance of vOTUs in each sample was estimated as Transcripts per Million (TPM) and used to calculate Bray-Curtis dissimilarity between samples. A PERMANOVA analysis and Non-metric Multidimensional Scaling (NMDS) of viral abundance reveals a grouping of viral communities according to plant compartment and phenotype, consistent with a potential influence of plant phenotype in viral community composition (Figure 1d). Interestingly, the gradient across compartments follows a pattern of proximity, with the rhizosphere positioned between the bulk sediment and root, consistent with their spatial arrangement relative to the plant.

### 2.2 Viral Population Co-occurrence Suggests Influence of Plant Phenotypes and Compartments on Viral Community Composition

We examined how vOTUs were distributed across different plant phenotypes and compartments, focusing on viral sequences found in more than one sample (black bars in Figure 2a), and compared vOTU distributions to a null model randomized while conserving only richness per phenotype and compartment (differences reported are based on *χ*^2^ tests at a 0.05 significance level). Of the 769 vOTUs identified in this study, almost half (42%) were found in only one sample (Figure 2a), suggesting that viral sequences were not widely recovered. A few vOTUs are cosmopolitan, with two particular sequences appearing in 19 of the 24 samples (we were unable to taxonomically characterize these vOTUs; see Methods for details on taxonomic characterization). Of the 444 vOTUs found in multiple samples, only 62 (18%) were present in samples from different phenotypes (Figure 2b). This is lower than expected by the null model and indicates that viral populations are less shared between short and tall plants than we would expect by chance, highlighting a strong influence of plant phenotype on viral community composition.

**Figure 2:**
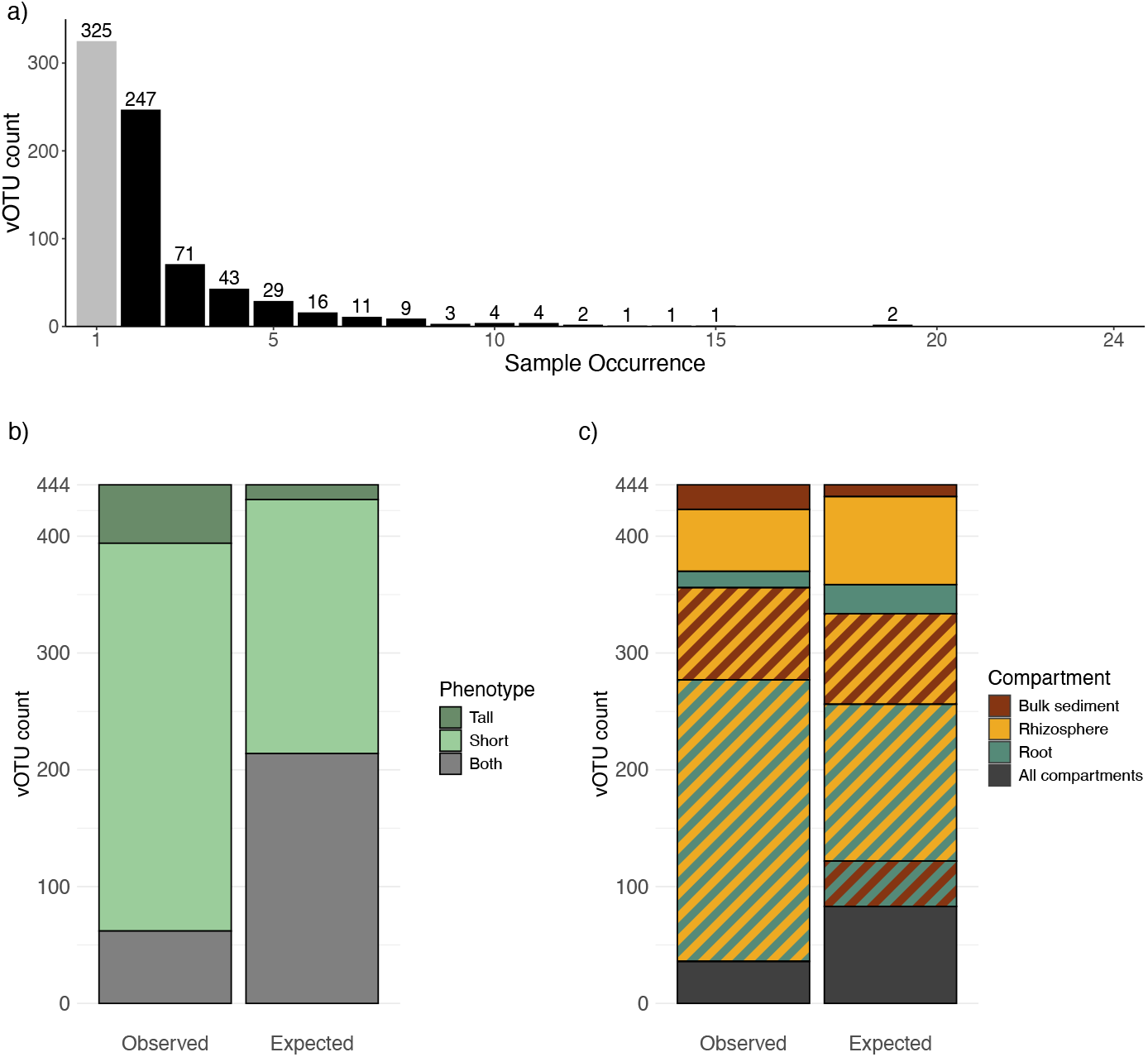
vOTU co-occurrence across phenotypes and compartments. (**a)** Number of samples in which a single vOTU is found. Most vOTUs (444 out of 769) are found in 2 or more samples (black bars), with two sequences found in 19 out of the 24 samples. **(b)** Comparison of plant phenotypes associated with vOTUs that appear in two or more samples (observed data) to a null model of co-ocurrence (expected). Less vOTUs are found in samples obtained from plants with different phenotypes (gray section in left bar) than expected (gray section in right bar). Statistical differences based on *χ*^2^ test (p-value *<* 0.05). (**c)** Comparison of microbiome compartments associated with vOTUs that appear in two or more samples (observed data) to a null model of co-occurrence (expected). Fewer vOTUs are found in samples from all three soil compartments (gray section in the left bar) than expected (gray section in the right bar). More vOTUs are found in samples from two different compartments than expected (stripe sections), with the largest increase in vOTUs shared between the rhizosphere and root. No vOTUs are found to be shared exclusively between the bulk and root compartments. Fewer vOTUs are found in samples from the same compartment than expected (solid colors). Statistical differences based on *χ*^2^ test (p-value *<* 0.05)

When we examined vOTU distribution across compartments, we found that 88 out of 444 (20%) were only found in one compartment (bulk sediment, rhizosphere, or root). Meanwhile, 320 vOTUs (72%) appeared in two compartments, and 36 vOTUs (8%) were present in samples from all three compartments. Contrary to the phenotype pattern, we observed more vOTU co-occurrence between compartments than expected by chance. This was mainly driven by many vOTUs being shared between two compartments. Among vOTUs found in only one compartment, we observed more vOTUs exclusive to the bulk sediment than expected, while fewer were exclusive to the rhizosphere or root.

Interestingly, while 36 vOTUs were found in all three compartments, none were shared exclusively between the bulk sediment and the root. All vOTUs found in both compartments were also present in the rhizosphere. Similarly, there was a lower proportion of vOTUs shared between the bulk sediment and rhizosphere than expected, but a higher proportion shared between the rhizosphere and root.

Overall, we found that vOTUs are less shared across plant phenotypes than expected, but more shared across compartments, especially those that are spatially close. These results suggest that both plant phenotype and compartment are key factors shaping viral community composition.

### 2.3 Taxonomic characterization of viral populations showcases novelty and similarities with aquatic and soil associated viral populations

We used vContact2 [21] to assign viral taxonomy to the newly identified vOTUs. vContact2 uses cluster protein similarity to predict genus-level taxonomy. Subclusters in the vContact2 output represent genus-level similarity, meaning that if two sequences are in the same subcluster they are expected to belong to the same genus. The 769 vOTUs identified in this study were clustered along with viral RefSeq, and vOTUs characterized from aquatic and terrestrial habitats, recovered from the PIGEON [14] and GOV [19] databases (Figure 3). Of the 769 vOTUs identified, only 139 (18%) match viral sequences previously described, forming 102 distinct clusters with viral sequences derived from other metagenomic studies sampling diverse soil and aquatic systems. None of the vOTUs from this study clustered with RefSeq sequences. The remaining 630 vOTUs are either singletons (348 in total) or part of 106 clusters (282 vOTUs), all of which contain only sequences from this study.

**Figure 3:**
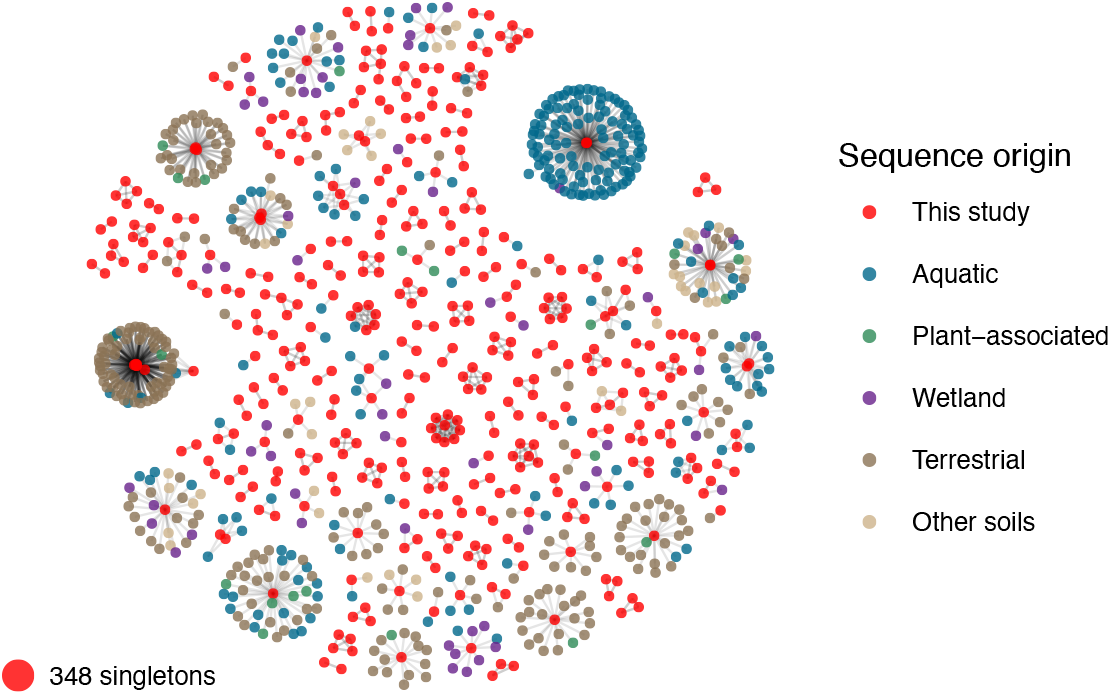
Taxonomic characterization of viral populations. Viral taxonomy of the vOTUs was predicted using vContact2. vOTUs in the same cluster are expected to share genus-level similarities. Of the 769 vOTUs, 421 were assigned to 207 clusters, while 348 were classified as singletons. Among the clustered vOTUs, 102 clusters containing 139 vOTUs matched viral populations previously reported in other environmental studies. The network displayed here represents viral clusters, with edges indicating only the direct similarities to the vOTUs identified in this study. Other potential interactions among vOTUs not described in this study are not depicted.

Most of the vOTUs characterized here did not share significant protein cluster similarity with known viruses and are therefore considered novel. This highlights the vast diversity of viral species across the globe and underscores how little we have yet managed to characterize the full extent of viral diversity.

### 2.4 Viral populations infect hosts that mediate biogeochemical cycles

We next set out to determine what microbial hosts are being targeted by the viral populations found in cordgrass-associated metagenomes. In prior work, bacterial populations found in *S. alterniflora* associated microbiomes were shown to possess nitrifying, iron-oxidizing, sulfur-oxidizing, and sulfate-reducing properties [4]. We used iPHoP (integrated Phage Host Prediction) to predict host taxonomy for our vOTUs [22]. Singular host predictions were assigned to 64 out of 769 vOTUs (8.3%). The predicted hosts comprise 14 unique phyla and 22 orders. While iPHoP can predict host genus level, taxonomic levels lower than phylum are sometimes unknown. For example, 12 vOTUs with phylum level predictions had unknown order classifications.

The percentage of vOTUs with host predictions varied between 6% and 27% across different compartment and phenotype combinations (Figure 4a). Host richness was higher in the short phenotype across all compartments, consistent with the greater vOTU richness observed in viral communities associated with short phenotypes compared to tall ones. *Pseudomonadota* and *Thermodesulfobacteriota* comprise the largest proportion of targeted hosts for all compartments and phenotypes.

**Figure 4:**
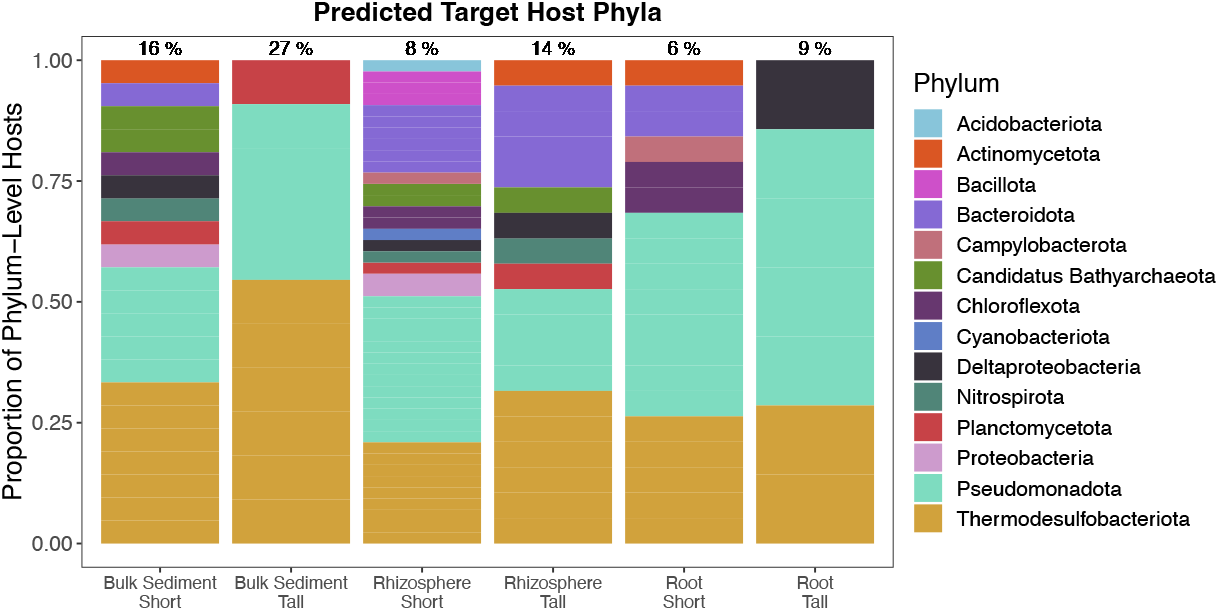
Host prediction reveals viral targeting of microorganisms implicated on important biogeochemical cycles in salt marshes. Hosts were predicted for the vOTUs associated *S. alterniflora*. 55 unique hosts were predicted at the phylum level for 64 vOTUs. Percentages above bars represent the proportion of vOTUs for which hosts were predicted out of all vOTUs found per phenotype and compartment combination. The absolute number of vOTUs with hosts predicted for each sample type were 21, 11, 43, 19, 19, and 7 for short bulk sediment, tall bulk sediment, short rhizosphere, tall rhizosphere, short root, and tall root, respectively. Each identified phylum is represented by a different color.

Host order-level taxonomy was used to infer the potential biogeochemical properties of the hosts targeted by viruses associated with *S. alterniflora*. Of the 22 unique host orders targeted by the vOTUs, 9 include genera capable of sulfur oxidation, sulfate reduction, and iron oxidation [4] (Suppl. Table 1 and 2). Half of the vOTUs with predicted hosts target taxa linked to at least one of these biogeochemical properties associated with nutrient cycling. However, no hosts were matched to known nitrifying bacteria. We also searched for Auxiliary Metabolic Genes (AMGs) in the viral sequences but did not identify any, based on our search criteria (see Methods).

## 3 Discussion

We investigated the viral populations associated with the *S. alterniflora* rhizosphere across three distinct microbiome compartments—bulk sediment, rhizosphere, and root—and two plant phenotypes, tall and short. We used a viromics pipeline to identify 769 putative viral sequences across all 24 metagenomic samples, with the vast majority being novel at the genus level. Our analysis points to niche differentiation as both plant phenotype and microbiome compartment are significant factors influencing viral community diversity and composition. Specifically, viral communities are more diverse when associated with short phenotype plants across most microbiome compartments, with the rhizosphere of short plants being the most diverse. Viral communities differ markedly across plant phenotypes, but viral populations are shared among sediments, especially between rhizosphere and root. We predicted virus-host interactions and found that viral populations associated with *S. alterniflora* target microbial groups potentially involved in sulfur oxidation, sulfate reduction, and iron oxidation. To our knowledge, this is the first study examining viral community composition in the root zone of foundational wetland plants.

Observed differences in viral diversity across plant phenotypes and compartments contrast with previous findings on microbial communities. While we detected increased viral diversity in the rhizosphere and root compartments of short phenotype plants, no corresponding differences were observed in microbial diversity in these same compartments [4]. Similarly, although viral diversity varied between the rhizosphere and bulk sediment of short plants, microbial diversity showed no significant difference between these compartments. These results suggest that viral diversity, as inferred through analysis of viral sequences in bulk metagenomes, does not necessarily align with microbial diversity, underscoring the complexity of microbe-virus interactions.

Viral community composition varied strongly between plant phenotypes. We hypothesize that this variation reflects differences in microbial composition between short and tall plants, as microbial community composition has been shown to influence viral communities [11, 13]. Near the creek bank of Sapelo Island, Georgia, the sediments are more oxidized and there is a faster turnover of nutrients mediated by microbial activity, while inland sediments are more stressful for plants due to increased sulfate reduction and salt concentration, influencing plant traits [3, 4, 17, 23]. As a result, plants located closer to the tidal banks exhibit taller phenotypes, while those farther inland show shorter phenotypes [24]. Microbial communities differ across phenotypes and are enriched with taxa known to catalyze distinct biogeochemical processes, which may reflect the physiological stress associated with the phenotypic gradient [4]. Additionally, differences in viral communities between phenotypes may be influenced by soil physicochemical properties, as shown in other wetlands [13], and the spatial distances between plants, as viral communities in soils exhibit high spatial heterogeneity [25].

Taxonomic classification of the viral populations revealed that most viral sequences are novel at the genus level, although some are related to genera previously identified in metagenomic studies of aquatic and soil-associated viral communities. This result highlights the unexplored viral diversity in marine sediments. Based on our prediction of virus-host interactions, we contend that the viral populations associated with the *S. alterniflora* root zone target microbial groups potentially involved in sulfur oxidation, sulfate reduction, and iron oxidation. These microbes are abundant in this environment [4], suggesting that viruses may be influencing key biogeochemical processes and overall ecosystem dynamics.

A key limitation of our study, highlighted by the rarefaction curve in Figure 1b, is that our sampling does not fully capture the diversity of viral communities. This may be partially due to the use of bulk metagenomes. Enriching samples by isolating VLPs has been shown to be more effective in the recovery of viral populations found in complex samples such as soils or marine sediments [26]. Moreover, our DNA-based study overlooks RNA viruses, which are often underrepresented in metagenomic studies. Additionally, the relatively short length of our viral sequences (with a minimum of 5kb) implies that many vOTUs are fragments of viral genomes. Future studies focusing on viromes rather than bulk metagenomes, could offer a more complete picture of the *S. alterniflora* associated viral communities. Similarly, assembly issues may have biased viral sequence recovery, as highly diverse metagenomes are often harder to assemble. The higher viral richness observed in short plants could result from greater microbial diversity within tall plants, leading to poorer assembly and thus fewer vOTUs detected. To address this, we resorted to various assembly strategies and conducted multiple rounds of viral predictions, as described in previous studies [27]. In addition, our ability to make robust conclusions regarding host predictions is limited by the fact that we were only able to predict hosts for a small proportion of the vOTUs. While this is a common challenge in metagenomic studies, it limits the explanatory power of our findings.

In conclusion, our study provides insights into the viral populations associated with the *S. alterniflora* root zone across different microbiome compartments and plant phenotypes. While we identified a diverse range of putative viral sequences, the limitations of our sampling methods—such as the use of bulk metagenomes and the challenges of assembling highly diverse metagenomic data—mean that our analysis offers only a partial view of the diversity, metabolic potential, and ecological impacts of these viral communities. Despite these constraints, we observed significant differences in viral richness and community composition across plant phenotypes and compartments, suggesting niche differentiation of viral populations with plant traits and proximity to plant roots serving as forces in shaping the viral communities. To gain a more comprehensive understanding of the complex interactions between viruses, microbes, and plants, future studies should focus on isolating viral particles and incorporating RNA viruses, while quantifying the role of viruses in microbial community functioning of wetland environments. Ultimately, our findings highlight the unresolved diversity of viral communities in marine sedimentary ecosystems while underscoring the need for a more in-depth exploration of the viral diversity in coastal wetlands and their potential ecological significance.

## 4 Materials and Methods

### 4.1 Sample collection and processing

The sequencing data used in this study has previously been published in [4]. Briefly, 24 samples were recovered in Summer 2018 and 2019 from *S. alterniflora* associated salt marsh at Sapelo Island, Georgia, USA for subsequent microbial metagenomic analysis. The samples expand 4 replicates for each of the 6 combinations of two plant phenotypes (short and tall), and 3 microbiome compartments (bulk sediment, rhizosphere, and root). For DNA extractions, the rhizosphere and root microbiomes were separated using sonication in an epiphyte removal buffer, following the method described in [4, 28]. DNA extractions were performed using the DNeasy PowerSoil kit following the manufacturer’s instructions. Shotgun metagenome sequencing was performed on an Illumina NovaSeq 6000 S4 2×150 Illumina flow cell at the Georgia Tech Sequencing Core (Atlanta, GA).

### 4.2 Read trimming and assembly

We assessed the read quality of the Illumina paired-end DNA sequencing reads with FastQC v.0.11.9 before and after read trimming [29]. We obtained trimmed reads using a QC of 20 and adapter trimming in Trimmomatic v0.39. [30]. We used MEGAHIT v1.0.2 with default parameters to assemble [31]. Four assembly strategies were conducted to increase the number of assembled contigs. First, we assembled the reads of each of the 24 individual samples. Second, 6 coassemblies were conducted by concatenating all reads in the 4 replicates grouped by soil compartment and plant height. Then, we mapped sample reads to the *S. alterniflora* genome (GeneBank ID GCA 008808055.2) using BWA v0.7.1 [32]. Samtools v1.16 [33] was used to extract unmapped reads. We repeated the individual assembly and co-assembly strategies using the unmapped reads. Our strategy yielded a total of 60 assemblies.

### 4.3 Viral population identification

We used VirSorter2 v2.2.1 [18] with default parameters to detect virus-like fragments from the MEGAHIT assembled contigs. Contigs shorter than 5,000 bp were removed from the viral sequence prediction pool. CD-HIT v4.5.4 [34] was used to cluster the sequences into viral populations based on sequence similarity with 95% average nucleotide identity matches and 80% query coverage. These cutoffs have been previously shown to define dsDNA viral populations into discrete genotypic groups [19]. One representative sequence was kept for each viral population.

We performed viral-specific reassembly to increase the number of viral populations by following the pipeline used in [27]. We mapped trimmed reads to the 1280 putative viral populations using BWA. The reads that mapped with any amount of coverage were co-assembled with MEGAHIT. Viral sequences were predicted from these assemblies using VirSorter2 with a 1,500 bp minimum length. Predicted viral sequences with a length shorter than 5 kb were removed from the pool. These sequences were added back to the original viral populations and clustered using CD-HIT with the cut-offs previously discussed. CheckV [35] with default database and parameters was used to filter the putative viral populations predictions for quality and completeness. Sequences that did not contain a viral-like protein, according to CheckV, were removed from further analysis.

Raw reads were mapped to the PIGEON and GOV 2.0 databases. The PIGEON database contains 266,125 viral population sequences from a variety of ecosystems including freshwater and marine sources, peatlands, and other soils [14]. The GOV 5kb database contains 488,130 sequences obtained from the Tara Oceans global oceanographic research expedition [19]. We assumed a sequence was present in our samples if there is a minimum 75% coverage. We pooled the database-recovered sequences with our *de novo* predicted viral populations. Duplicated sequences were removed and quality was checked with CheckV for the database-recovered sequences. This viral population collection was used for the analyses presented in this study.

Abundance and richness analysis were performed using read mapping. We mapped the per sample raw reads to the viral populations collection with BWA and calculated read coverage and depth with Samtools. As before, a viral population was assumed to be found in a given sample if the mapping coverage was higher than 75%. TPM (transcripts per kilobase million) was used as a measure of relative viral abundance.

### 4.4 Gene prediction and annotation for AMG identification

Gene prediction was done using Prodigal V2.6.3 [36] with default parameters. The annotated protein file was then used as input for gene annotation using HMMER V3.3.2 and the PFAM [37], Eggnog 10239 (viral) [38], and pVOG [39] databases. A bit score above 30 and an e-value cutoff of 0.01 were used to select significant hits. Genes were considered AMGs (Auxiliary Metabolic Genes) only if they were found in the center regions of the viral sequence, surrounded by annotated viral proteins.

### 4.5 Viral Taxonomy and Host identification

Viral taxonomy was inferred using vContact2 with default parameters [21]. Briefly, vContact2 clusters viral sequences based on protein similarities. Predicted subclusters are meant to represent viral taxonomy at the genus level. Our viral sequences were compared to viral sequences in Refseq 211, GOV [19] and a subset of the PIGEON database containing predicted viral sequences from plant and soil ecosystems [14].

iPHoP (Integrated Phage Host Prediction) v1.3.0 [22] was used to identify potential hosts infected by our viral populations using the default host database. The predict command was used with default parameters. Refseq and Genbank IDs from the iPHoP genus level predictions were used to confirm taxonomy. A list of microorganisms exhibiting sulfate-reducing, sulfur-oxidizing, nitrifying, and iron-oxidizing capabilities was retrieved from [3]. Given the challenge of identifying the precise microbial species a virus may infect, we compared the predicted viral hosts to microorganisms with relevant biogeochemical functions at the order level. The taxonomy of both predicted hosts and known prokaryotes with these properties was manually verified using the NCBI Taxonomy Database.

### 4.6 Data and code availability

Sequencing data was reanalyzed from [4]. The viral population sequences analyzed in this study, and all code are available at https://github.com/WeitzGroup/SaltMarshViral.git and archived at https://doi.org/10.5281/zenodo.15102500

Figures were created in R version 4.3.2 [40]. The stats version 4.3.2 package was used to perform wilcoxon and chi-squared tests in R. The R package vegan v2.6 was used for PERMANOVA test. Figure code for vOTUs co-occurrence and taxonomy depiction was adapted from [11] and [41], respectively.

### 4.7 Supplementary Material

Supplementary Table 1: Host prediction list

Supplementary Table 2: Biogeochemical properties of host orders

## 5 Acknowledgments

This work was supported by grants from the Simons Foundation Life Sciences Program (722153) and National Science Foundation (DEB-1934586) to JSW, and by an institutional grant (NA18OAR4170084) to the Georgia Sea Grant College Program from the National Sea Grant Office, National Oceanic and Atmospheric Administration, US Department of Commerce, by a grant from the National Science Foundation (DEB 1754756) and by the DOE Office of Science, Office of Biological and Environmental Research (BER; DE-SC0023297) to JEK. Funding sources had no role or influence on study design, analysis, interpretation, or submission. We thank Paula Saavedra for illustration contributions. This research was supported in part through research cyber infrastructure resources and services provided by the Partnership for an Advanced Computing Environment (PACE) at the Georgia Institute of Technology, Atlanta, Georgia, USA.

